# Germline genetic determination of cancer progression and survival

**DOI:** 10.1101/576181

**Authors:** Benjamin N. Ostendorf, Kimia N. Tafreshian, Nneoma Adaku, Jana Bilanovic, Bernardo Tavora, Sohail F. Tavazoie

## Abstract

We report the surprising finding that common germline polymorphisms of *APOE*, present in approximately 39% of Caucasians, predict survival outcomes in human melanoma. Analysis of The Cancer Genome Atlas revealed that carriers of the *APOE2* variant experienced shorter survival relative to *APOE3* homozygotes, while *APOE4* variant carriers exhibited increased survival. Consistent with this, melanoma growth in human *APOE* knock-in mice followed the order of *APOE2* > *APOE3* > *APOE4*, revealing causal regulation of progression by *APOE* variants. Mechanistically, recombinant ApoE protein variants differentially suppressed melanoma cell invasion and endothelial recruitment phenotypes. Moreover, tumors in *APOE4* mice exhibited greater immune cell infiltration and activation relative to tumors of *APOE2* mice. These findings support the notion that human germline genetic makeup can impact the trajectory of a future malignancy.

---

The secreted glycoprotein Apolipoprotein E (ApoE) was previously found to suppress melanoma progression and metastasis through cancer cell autonomous and non-autonomous mechanisms (1–3). In humans there are three prevalent ApoE variants termed ApoE2, ApoE3, and ApoE4, which differ in only two amino acids and exhibit differential binding to ApoE receptors (Fig. 1 a)(4–7). The *APOE4* variant is the strongest monogenetic risk factor for Alzheimer’s disease, while *APOE2* confers reduced risk for this disease (8, 9). *APOE* variants also modulate the risk for cardiovascular disease (reviewed in (10)). Given their high prevalence in the general population, the association of *APOE* variants with a number of human phenotypic outcomes has been investigated. However, a causal role for these variants in disease pathogenesis has only been demonstrated for Alzheimer’s disease and atherosclerosis through the use of murine models (11, 12). The potential association between *APOE* status and cancer outcomes has remained inconclusive (13, 14).

**Figure 1.**
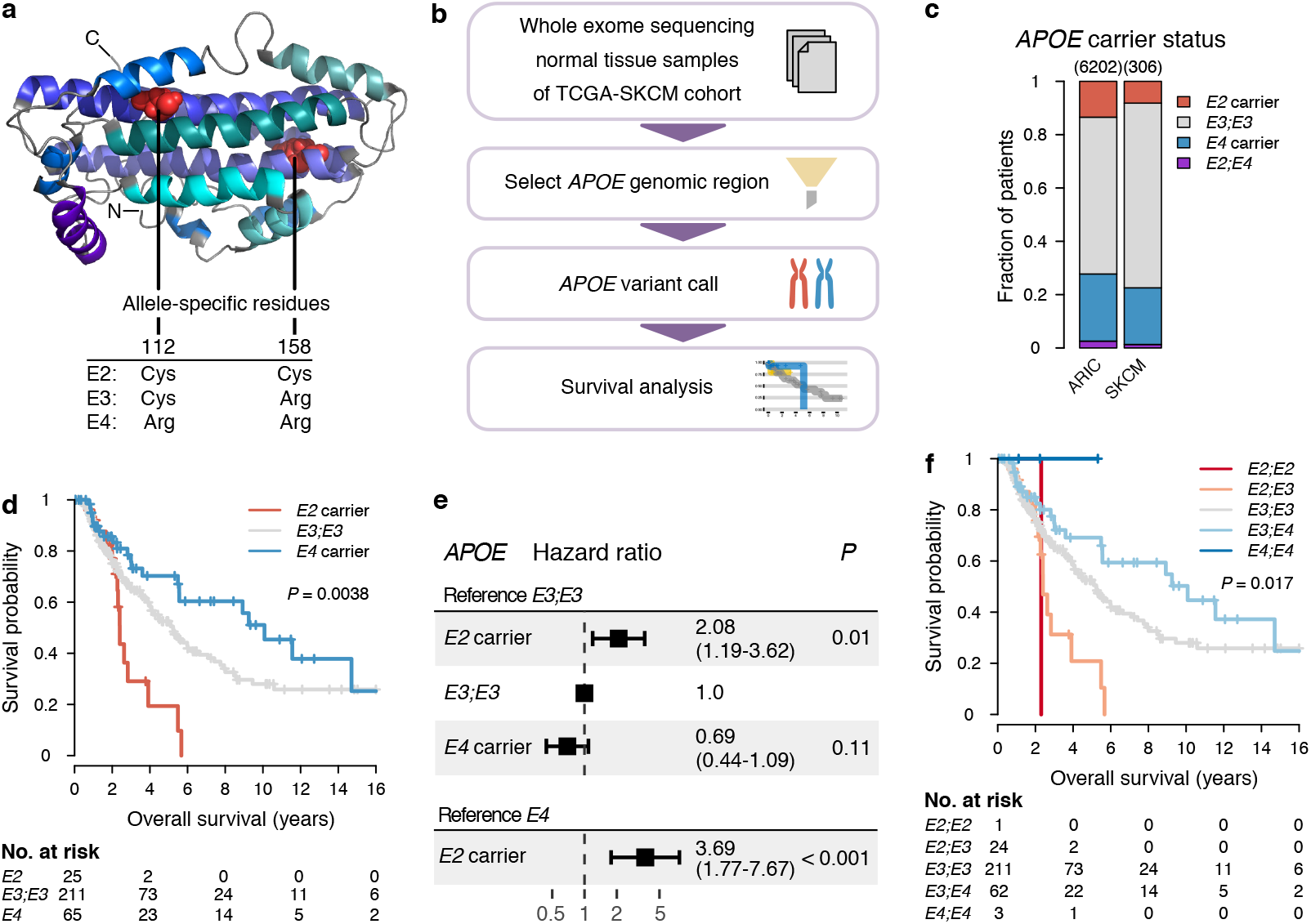
*APOE* germline variants predict survival in human melanoma. **a**, Structural representation of ApoE3 (Chen et al., PNAS, 2011). **b,** Computational pipeline to analyze the impact of *APOE* genotype on melanoma outcome. **c,** Proportion of *E2* and *E4* carriers in the *Atherosclerosis Risk in Communities* study (ARIC) and in patients with stage II/III melanoma in the TCGA-SKCM cohort (*P* = 0.0017, χ^2^ test). **d-e,** Survival **(d)** and hazard ratios **(e)** of stage II/III melanoma patients stratified by *APOE* carrier status (p-values according to log-rank test (d) or a Cox proportional hazards model (e); numbers in parentheses indicate 95% confidence interval). **f,** Survival of stage II/III melanoma patients stratified by bi-allelic *APOE* genotype (logrank test).

Given prior evidence supporting a role for ApoE in melanoma progression, we investigated the association between germline *APOE* variant status and clinical outcome of patients with advanced localized melanoma (stages II-III) by determining *APOE* genotypes from normal tissue whole exome sequencing data of participants in the TCGA-SKCM study (Fig. 1b) (15). In comparison to a control group with similar age and ethnic composition (16), homozygotes for the most common *APOE3* variant (i.e., non-*E2/E4* carriers) were modestly overrepresented in the TCGA-SKCM cohort, indicating that neither *APOE2* nor *APOE4* carrier status predisposed for increased melanoma incidence (Fig. 1c). Strikingly, however, *APOE* carrier status was significantly associated with survival (median survival of 2.4, 5.2, and 10.1 years for *E2* carriers, *E3;E3* homozygotes, and *E4* carriers, respectively; p = 0.0038, log-rank test; Fig. 1d). Cox proportional hazard regression analysis revealed an increased hazard ratio for *E2* carriers versus *E3;E3* homozygotes (HR = 2.08, p = 0.01) and versus *E4* carriers (HR = 3.69, p < 0.001). There were no significant differences in potentially confounding characteristics such as gender, age, tumor stage, Clark level, Breslow depth, and mutational or transcriptomic subtype between *APOE* carrier groups at the time of diagnosis (Extended Data Fig. 1a-g). Stratifying patients by bi-allelic *APOE* genotype was consistent with a gene dosage effect on survival (median survival of 2.3, 2.4, 5.2, 10.1 years and median survival not reached for *E2;E2, E2;E3, E3;E3, E3;E4*, and *E4;E4*, respectively; p = 0.017, log-rank test) (Fig. 1f and Extended Data Fig. 1h). Thus, germline genetic variants of *APOE* genotype were significantly associated with survival in patients with advanced localized melanoma.

In order to assess whether *APOE* genotype merely correlated with or was causally involved in altering melanoma outcome, we used mice in which the endogenous murine *Apoe* locus is replaced with one of the three human *APOE* variants (11, 17, 18). Using the YUMM1.7 murine melanoma model, we observed significantly slower tumor progression in *E4;E4* and faster tumor progression in *E2;E2* mice relative to *E3;E3* mice, consistent with *APOE* genotype causally impacting melanoma progression in a manner analogous to the observed clinical association (Fig. 2a). In order to assess the impact of *APOE* genotype on metastatic progression of melanoma we used the aggressive B16F10 mouse melanoma model which exhibits efficient lung colonization upon intravenous injection. *APOE2* mice exhibited a significantly enhanced metastatic progression rate relative to *APOE3* and *APOE4* mice (Fig. 2b).

**Figure 2.**
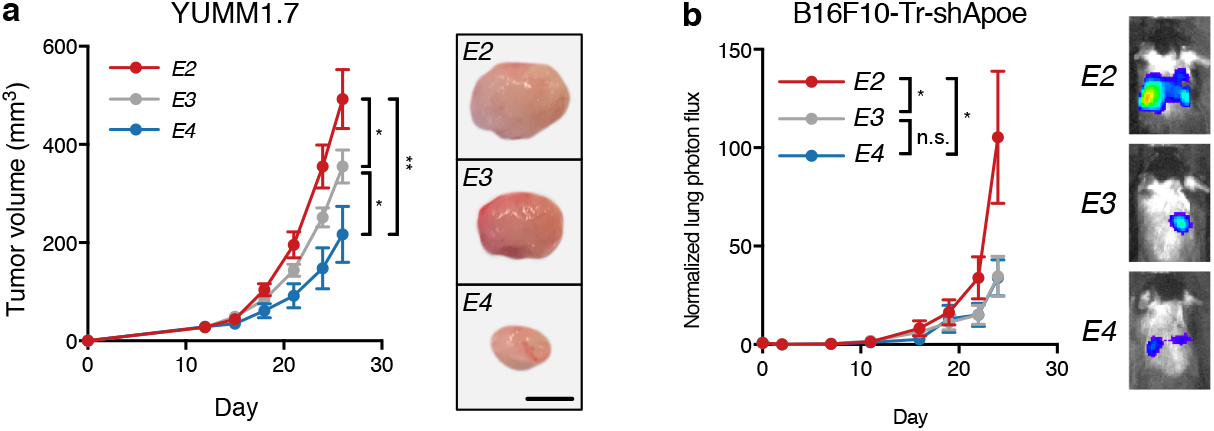
Human ***APOE*** variants modulate progression in mouse melanoma models. **a**, Tumor growth of 1 × 10^5^ YUMM1.7 cells in *APOE* knock-in mice (n = 9-11 per group, one-tailed t-test; representative of two independent experiments). Representative tumors correspond to day 26 (scale bar, 3 mm). **b**, Bioluminescence imaging of metastatic progression of 1 × 105 B16F10-TR-shApoe cells intravenously injected into *APOE* knock-in mice (n = 10 per group; one-tailed Mann-Whitney test; representative of two independent experiments). Images correspond to representative mice on day 24 after injection. \**P* < 0.05, \*\**P* < 0.01.

Our findings in primary tumor growth and metastatic colonization models of melanoma revealed enhanced progression in *APOE2* mice relative to *APOE3 and APOE4* mice, consistent with the clinical association observations. The differential impact of *APOE3 and APOE4* on progression, however, was only observed in the aforementioned primary tumor growth assay but not in the metastatic colonization assays. To confirm that *APOE3 and APOE4* differentially impact melanoma progression, we employed a second independent model of melanoma progression. We hypothesized that a genetically induced model of melanomagenesis might provide better resolution and sensitivity for capturing a potential difference between these variants. To this end, we crossed *APOE3* and *APOE4* mice to *Tyr::CreER;Braf^V600E/+;^Pten^lox/lox^* mice, in which melanoma formation can be initiated through Tamoxifen-induced and Cre-mediated excision of *Pten* in melanocytes (19) (Fig. 3a). This five gene loci and one transgene compound mutant expressed ApoE variants in both the tumor and host compartments. Mice with the *E4;E4* genotype exhibited reduced melanoma tumor burden in comparison to *E3;E3* mice, consistent with our results in the transplantable melanoma tumor progression model (Fig. 3b). These findings reveal that human *APOE* alleles differentially regulated progression in murine melanoma models, consistent with the clinical association data described above.

**Figure 3.**
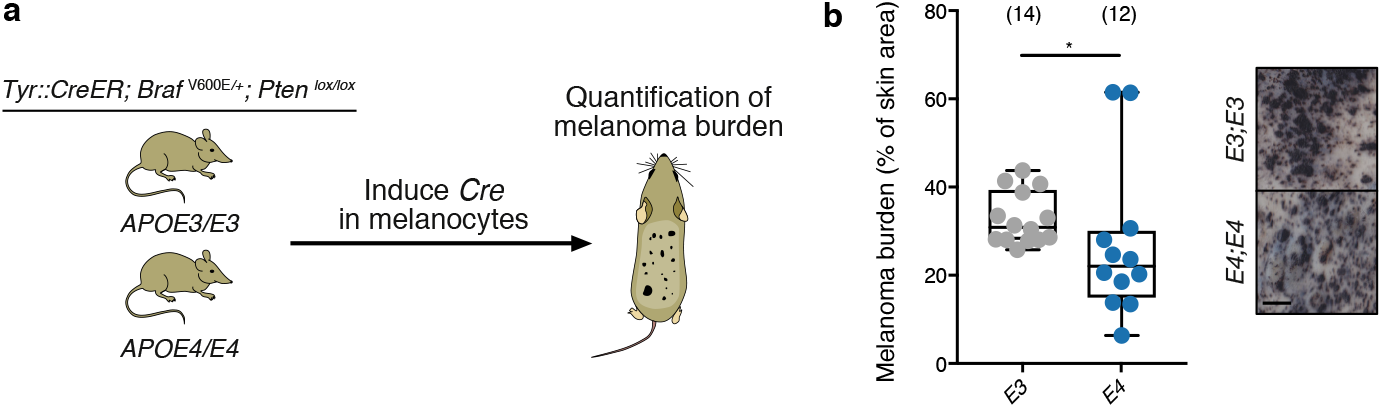
Human *APOE* variants impact progression in a genetically induced melanoma model. **a,** Experimental approach to assess the impact of human *APOE* alleles on genetically induced mouse melanoma progression. **b,** Melanoma burden of mice described in (a) on day 35 after induction (*P* = 0.01, one-tailed Mann-Whitney test). Images of representative skins are shown on the right. Circles correspond to individual mice and numbers in parentheses indicate sample sizes. \**P* <0.05.

ApoE exerts pleiotropic anti-tumor effects by suppressing melanoma cell invasion and endothelial recruitment (2). We thus assessed whether ApoE variants differentially affect these phenotypes. Recombinant ApoE3 and ApoE4 proteins were more efficient than ApoE2 in suppressing Matrigel invasion of B16F10 melanoma cells depleted of endogenous ApoE (Fig. 4a). ApoE3 and ApoE4 were also more potent than ApoE2 in suppressing endothelial recruitment by highly metastatic human melanoma MeWo-LM2 cells (Fig. 4b). These findings reveal that the ApoE2 variant, which confers worse prognosis in humans and enhanced melanoma progression in mice, is less effective at suppressing multiple cancer progression phenotypes.

**Figure 4.**
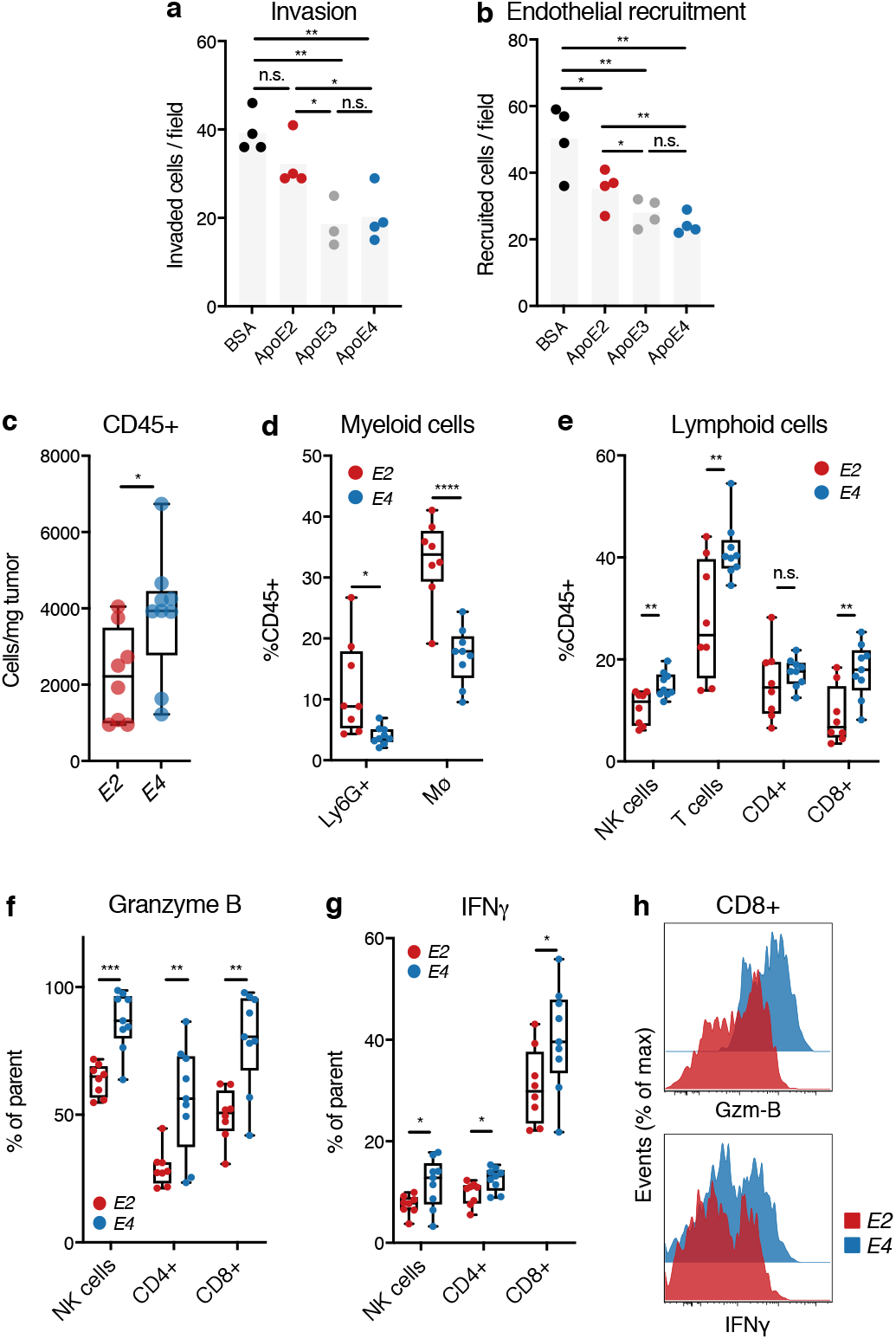
Human *APOE* variants modulate key tumor progression phenotypes. **a**, Matrigel invasion by 1 × 10^5^ B16F10-TR-shApoe cells treated with the indicated recombinant proteins (n = 3 – 4; one tailed t-test). **b,** Trans-well recruitment of 1 × 10^5^ human umbilical vein endothelial cells treated with the indicated recombinant proteins by 5 × 10^4^ human melanoma MeWo-LM2 cells (n = 4; one tailed t-test). **c,** Number of CD45+ leukocytes per tumor weight infiltrating YUMM1.7 tumors in *APOE2* or *APOE4* knock-in mice (P = 0.02, one-tailed t-test). **d-e,** Proportion of myeloid **(d)** and lymphoid **(e)** immune cells in YuMM1. 7 melanoma-bearing *APOE2* and *APOE4* mice (two-tailed t-tests). **f-h,** Expression of Granzyme B (Gzm-B) **(f)** and Interferon-γ (IFNγ) **(g)** in immune effector cells infiltrating YUMM1.7 melanomas in *APOE2* versus *APOE4* mice (one-tailed t-test). **h,** Representative flow cytometry plots illustrating the expression of activation markers in YUMM1.7-infiltrating CD8+ T cells in *APOE2* and *APOE4* mice. Data are representative of three (ab) or two (c-h) independent experiments. Circles in (c-g) correspond to individual mice. \**P* < 0.05, \*\**P* < 0.01, \*\*\**P* < 0.001, \*\*\*\**P* < 0.0001. n.s., nonsignificant.

ApoE was previously shown to also enhance anti-tumor immunity by repressing the survival and abundance of an innate immunosuppressive immune cell population termed granulocytic-myeloid derived suppressor cells (G-MDSCs, also referred to as polymorphonuclear MDSCs) (1). To determine whether *APOE* genotypes differentially impact the immune response in cancer, we assessed the tumor immune microenvironment in *APOE2* and *APOE4* mice *in-vivo* by flow-cytometry. *APOE4* mice exhibited enhanced recruitment of CD45+ leukocytes into YUMM1.7 melanoma tumors relative to *APOE2* mice (Fig. 4c, Extended Data Fig. 2a-b). Within the CD45+ leukocyte compartment we observed a significantly reduced proportion of immunosuppressive Ly6G+ G-MDSCs in *APOE4* relative to *APOE2* mice (Fig. 4d). Concomitant with the reduction of this immunosuppressive population, we observed an increase in anti-tumor effector cell subsets in *APOE4* mice, such as natural killer (NK) and CD8+ T cells (Fig. 4e, Extended Data Fig. 4c-d). We also observed a reduction in tumor associated macrophages in *APOE4* mice (Fig. 4d). Intracellular flow cytometry revealed enhanced activation of immune effector cells in *APOE4* mice relative to *APOE2* mice, as illustrated by increased Granzyme B and Interferon-γ expression in NK, CD8+ and CD4+ T cells (Fig. 4f-h). These findings suggest that the differential impact of ApoE variants on melanoma progression likely results from alterations in multiple cell autonomous and non-cell autonomous phenotypes that collectively contribute to melanoma progression.

In summary, we show that highly prevalent variants of *APOE* exhibit differential effects on melanoma progression in mice and humans. These effects at the organismal level are accompanied by differential effects of ApoE variant proteins on cellular phenotypes affecting melanoma progression, including invasion, endothelial recruitment, and immune cell infiltration and activation. Our findings reveal ApoE2 to be hypomorphic and ApoE4 to be hypermorphic in suppressing cancer progression phenotypes. What is the biochemical basis for this difference? Our past work has shown that ApoE proteins mediate suppressive effects on melanoma cell invasion, endothelial recruitment, and G-MDSC survival by acting on ApoE receptors LRP1 and LRP8 (1, 2). ApoE variants have been previously shown to exhibit differential binding affinities to the ApoE receptor LDLR (E2 << E3 < E4) (5). Differential binding or signaling by distinct ApoE variants has also been reported for additional ApoE receptors such as LRP1 and LRP8 (6, 7, 20). Thus, the differential phenotypic effects of these variants on suppressing melanoma progression and prometastatic cellular phenotypes is consistent with their established receptor binding capacities for ApoE receptors. Our findings reveal the surprising observation that the *APOE4* variant, which is associated with increased risk of Alzheimer’s disease and atherosclerosis, is protective against melanoma progression. These findings have important clinical implications as they may help to identify patients at high risk for metastatic progression for treatment with systemic therapy. More generally, our findings reveal that germline genetic variants can impact progression of a common human cancer and raise the possibility of the existence of additional common gene variants that predict and govern progression outcomes in other common malignancies.

## Methods

### Impact of *APOE* genotype on survival in human melanoma in the TCGA-SKCM cohort

To assess *APOE* genotypes of patients with melanoma, we downloaded aligned whole exome sequencing *BAM* files sliced for the genomic coordinates chr19:44904748-44910394 (GRCh38) from the TCGA-SKCM project using the Genomic Data Commons API (15). We called *APOE* variants using the samtools/bcftools package (21), providing allele frequencies for chr19:44908684 (rs429358) and chr19:44908822 (rs7412) as determined in the Atherosclerosis Risk in Communities (ARIC) study (16) as a prior distribution. Normal tissue samples (either blood, solid tissue, or buccal cells) were available for 470 patients. No genotype could be determined in 10 patients. Additionally, patients that exhibited the *E2;E4* genotype (n = 5) were excluded from analyses except for genotype frequency assessment. *APOE* genotype abundance in the normal population was based on the assessment of Caucasian patients in the ARIC study (16).

Clinical data including survival times and clinical response were used as recently curated (22). The R package ‘TCGAbiolinks’ was used to add clinical data for Breslow depth and Clark level (23). To assess the impact of *APOE* genotypes on survival, Kaplan-Meier survival analysis was performed, and statistical significance was assessed with the log-rank test using the ‘survival’ and ‘survminer’ packages (24, 25). Hazard ratios were calculated according to a Cox proportional hazards regression model using the ‘survival’ R package (24). For visualization purposes, survival data were truncated at 16 years, after which there were only individuals in the *E3;E3* group remaining. All analyses were performed using R v3.5 (The R Foundation for Statistical Computing) and RStudio v1.1.3.

### Animal studies

All animal experiments were conducted in accordance with a protocol approved by the Institutional Animal Care and Use Committee at The Rockefeller University. Human *APOE2* (strain #1547), *APOE3* (#1548) and *APOE4* (#1549) targeted replacement (knock-in) mice were obtained from Taconic Biosciences. *Tyr::CreER;Braf^V600E/+^;Pten^lox/lox^* mice (19) were obtained from Jackson laboratories (#013590).

### Tumor growth studies

To assess the impact of *APOE* genotype on the growth of syngeneic melanoma we subcutaneously injected 1 × 10^5^ (YUMM1.7) cells into the flank of 7-10-week-old and sex-matched human *APOE* targeted replacement mice. Cells were injected in a total volume of 100 *μ*L and YUMM1.7 cells were mixed 1:1 with growth factor reduced Matrigel (356231, Corning) before injection. Tumor size was measured on the indicated days using digital calipers and calculating tumor volume as (small diameter)^2^ × (large diameter) × Π / 6.

### Tail-vein metastasis assays

For tail-vein assays, B16F10 cells stably expressing a retroviral construct encoding luciferase were used in order to assess cancer progression by bioluminescence imaging as described previously (3, 26). To assess whether *APOE* genotype impacts metastatic progression, 1 × 10^5^ cells resuspended in 100 *μ*L of PBS were injected into the tail vein of 6-8-week-old male human *APOE* knock-in mice. Bioluminescence imaging was performed approximately twice a week and the signal was normalized to the signal obtained on day 0.

### Genetically initiated model of melanoma progression

Human *APOE* targeted replacement (knock-in) mice were crossed with *Tyr::CreER;Braf^V600E/+^;Pten^lox/lox^* mice. To induce melanoma, 6-7-week-old female mice were injected intraperitoneally with 1 mg of Tamoxifen (T5648, Sigma-Aldrich) on three consecutive days. Tamoxifen solution was prepared by dissolving Tamoxifen powder in 100 % ethanol at 50°C for five minutes and subsequent tenfold dilution in peanut oil to yield a 10 mg/mL working solution. To assess melanoma burden, dorsal skin samples stretching from ears to hips were harvested on day 35 after induction, depilated with commercial depilation cream (Nair), washed with water and fixed in 4% PFA. Skins were then scanned and the percentage of pigmented melanoma lesion area was quantified using Cellprofiler v3 (27).

### Cell lines

Mouse B16F10 melanoma cells and human umbilical vein endothelial cells (HUVEC) were obtained from the American Tissue Type Collection and cultured according to the supplier’s conditions. B16F10 cells transduced with a retroviral construct to express luciferase (26) and shRNA targeting murine *Apoe* (shRNA clone TRCN0000011799; B16F10-TR-shApoe) were described previously (3). YUMM1.7 cells, originally derived from a *Braf^V600E/+^;Pten^-/-^;Cdkn2a^-/-^* mouse melanoma, were generously provided by Marcus Bosenberg (28). MeWo melanoma cells were originally obtained from the American Tissue Type Collection. The highly metastatic MeWo-LM2 subclone was described previously (2). B16F10 and MeWo cells were cultured in DMEM medium with Pyruvate and Glutamine (11995, Gibco) supplemented with 10 % FBS (F4135, Sigma), Penicillin-Streptomycin (15140, Gibco), and Amphotericin B (17-936E, Lonza). YUMM1.7 cells were cultured in DMEM/F-12 medium with L-Glutamine and 15mM HEPES (11330, Gibco) supplemented with 10 % FBS, Penicillin-Streptomycin, Amphotericin B, and 1 % non-essential amino acids (111400, Gibco). Contamination with mycoplasma was ruled out by PCR testing according to standard protocols (29).

### Mouse genotyping

Genotyping of *Tyr::CreER;Braf^V600E/+^;Pten^lox/lox^* mice was performed as recommended by the Jackson Laboratory. Genotyping to distinguish between mouse and human *APOE* was performed using standard PCR with independent reactions for mouse and human *APOE* (PCR product lengths of 200 bp and approximately 600 bp, respectively). In order to distinguish between human *APOE* alleles, we used PCR-based restriction fragment length polymorphism genotyping (30). In brief, a 244 bp fragment of *APOE* was amplified using standard PCR and digested with *HhaI* (R0139S, New England Biolabs), and allele-specific products were resolved on a 15% polyacrylamide gel. The following primers were used for the indicated PCR reactions:

*Tyr::CreER;Braf^V600E/+^;Pten^lox/lox^* mice

*Cre* transgene forward: 5’ – GCG GTC TGG CAG TAA AAA CTA TC – 3’
*Cre* transgene reverse: 5’ – GTG AAA CAG CAT TGC TGT CAC TT – 3’
*Cre* internal control forward: 5’ – CAC GTG GGC TCC AGC ATT – 3’
*Cre* internal control reverse: 5’ – TCA CCA GTC ATT TCT GCC TTT G – 3’
*Braf* forward: 5’ – TGA GTA TTT TTG TGG CAA CTG C – 3’
*Braf* reverse: 5’ – CTC TGC TGG GAA AGC GGC – 3’
*Pten*forward: 5’ – CAA GCA CTC TGC GAA CTG AG – 3’
*Pten* reverse: 5’ — AAG TTT TTG AAG GCA AGA TGC — 3’

Mouse versus human knock-in *APOE* mice

Common forward: 5’ – TAC CGG CTC AAC TAG GAA CCA T – 3’
Mouse *Apoe* reverse: 5’ – TTT AAT CGT CCT CCA TCC CTG C – 3’
Human *APOE* reverse: 5’ – GTT CCA TCT CAG TCC CAG TCTC – 3’

Human *APOE* allele restriction length polymorphism

Human *APOE* forward: 5’ – ACA GAA TTC GCC CCG GCC TGG TAC AC – 3’
Human *APOE* reverse: 5’ – TAA GCT TGG CAC GGC TGT CCA AGG A – 3’

### Isolation of tumor-infiltrating leukocytes

To isolate tumor-infiltrating leukocytes, YUMM1.7 tumors were resected on day 21 after injection and thoroughly minced on ice using scalpels. Tumor pieces were incubated in HBSS2+ (HBSS with Calcium and Magnesium (24020, Gibco) supplemented with 2% FBS, 1 mM sodium pyruvate (11360, Gibco), 25 mM HEPES (15630, Gibco), 500 U/mL Collagenase IV (LS004188, Worthington), 100 U/mL Collagenase I (LS004196, Worthington), and 0.2 mg/mL DNAse I (10104159001, Roche)) for 30 minutes at 37°C on an orbital shaker (80 rpm). After thorough trituration the mixture was passed through a 70 *μ*m strainer and diluted with HBSS2- (HBSS without Calcium and Magnesium (14170, Gibco), 2% FBS, 1 mM sodium pyruvate, and 25 mM HEPES). After centrifugation the cell pellet was resuspended in a 35 % Percoll solution (170891, GE Healthcare) and a phase of 70 % Percoll was underlaid using a glass Pasteur pipette. The resulting gradient was centrifuged at 800 × g for 20 minutes at room temperature without brakes. After removal of the red blood cell-containing pellet on the bottom and excess buffer containing cellular debris on the top, the cell population at the Percoll interphase enriched for tumor-infiltrating leukocytes was washed twice with HBSS2-.

### Flow cytometry

Unless otherwise mentioned all steps were performed on ice and under protection from light. Fc receptors were blocked by incubation with 2.5 *μ*g/mL anti-CD16/32 antibody (clone 93; 101320, BioLegend) in staining buffer (25 mM HEPES, 2 % FBS, 10 mM EDTA (351-027, Quality Biological), and 0.1 % sodiumazide (7144.8-16, Ricca) in PBS) for 10 minutes on ice. Cells were incubated with antibodies diluted in staining buffer for 20 minutes, washed with PBS, incubated with Zombie NIR Fixable Live/Dead Stain (423105, BioLegend) for 20 minutes at room temperature, and washed twice with staining buffer. Cells were analyzed on an LSR Fortessa (BD Biosciences). For cell quantification CountBright counting beads (C36950, Thermo Fisher) were added to the samples before analysis.

For intracellular staining of cytokines, cells were incubated with 500 ng/mL ionomycin (I0634, Sigma), 100 ng/mL Phorbol 12-myristate 13-acetate (P8139, Sigma), and 10 *μ*g/mL Brefeldin A (B7651, Sigma) for 3-4 hours at 37°C prior to surface labelling and live/dead staining as described above. Cells were then incubated in fixation/permeabilization buffer (00-5523, eBioscience) for 30 minutes on ice, washed with permeabilization buffer (00-5523, eBioscience), and incubated with antibodies diluted in permeabilization buffer for 20 minutes on ice. Finally, cells were washed with permeabilization buffer and subsequently with staining buffer.

### Antibodies

The following anti-mouse fluorophore-conjugated antibodies were used for flow cytometry: CD45 (clone 30-F11, BioLegend), B220 (clone RA3-6B2, BD Biosciences), CD11b (clone M1/70, BioLegend), Ly6G (clone 1A8, BD Biosciences), Ly6C (clone HK1.4, BioLegend), I-A/I-E (clone M5/11.15.2, BioLegend), F4/80 (clone BM8, BioLegend), CD24 (clone M1/69, BioLegend), CD103 (clone 2E7, BioLegend), CD19 (clone 1D3/CD19, BioLegend), TCRβ (clone H57-957, BioLegend), CD49b (clone HMa2, BioLegend), CD4 (clone GK1.5, BioLegend), CD8α (clone 53-6.7, BioLegend), Granzyme B (clone QA16A02, BioLegend), IFNγ (clone XMG1.2, BioLegend).

### Matrigel invasion assay

The assay was performed similarly to as described previously (2). B16F10-TR-shApoe mouse melanoma cells were serum starved overnight in DMEM supplemented with 0.2 % FBS. Prior to starting the assay, four Matrigel invasion chambers per condition (Corning, 354480) were equilibrated at 37°C with 500 *μ*L of 0.2% FBS DMEM in the top and bottom chambers. After 30 minutes, the starvation media in the top chamber was removed and replaced with 500 *μ*L of starvation media containing 1 × 10^5^ melanoma cells and either 10 *μ*g/mL of HEK293-produced recombinant ApoE2, E3, or E4 (Tonbo Biosciences 21-9195, 21-9189, and 21-9190), or an equimolar concentrations of BSA (20 *μ*g/mL; Sigma A3059). Chambers were then kept at 37°C for 24 hours to allow for invasion. Subsequently, the chambers were washed with 1× PBS, the tops were scraped with cotton swabs to remove residual non-invading cells, and the inserts were fixed in 4% paraformaldehyde for 20 minutes. After washing again with PBS, inserts were stained with DAPI (Roche, 10236276001) for 5 minutes, cut out, and then mounted bottom-up on slides with ProLong Gold Antifade Mountant (Invitrogen, P36930). Four representative images per insert were taken using a Zeiss Axiovert 40 CFL fluorescence microscope at 10× magnification, and the number of invaded cells was quantified.

### Endothelial recruitment assay

Human umbilical endothelial vein cells (HUVEC) were serum-starved overnight in EGM-2 media (Lonza, CC-3162) containing 0.2% FBS. Concurrently, 5 × 10^4^ highly metastatic MeWo-LM2 human melanoma cells were plated in a 24-well plate in DMEM supplemented with 10% FBS. On the day of the assay, the medium was replaced with EGM-2 starvation medium, and Mewo-LM2 cells were allowed to enrich the media for 6-8 hours at 37°C. Subsequently, BSA (20 ug/mL; Sigma, A3059) or ApoE2, E3, or E4 (10ug/ml; Tonbo Biosciences 21-9195, 21-9189, and 21-9190) were added to the media, and 3.0 *μ*m PET membrane inserts (Falcon, 353492) were placed in the wells. HUVEC cells were trypsinized, resuspended, and seeded equally into the top chambers. The cells were allowed to migrate for 16 to 18 hours, after which the inserts were mounted and analyzed as described for the invasion assay above.

### Statistical analysis

Unless otherwise noted all data are expressed as mean ± standard error of the mean. Groups were compared using tests for significance as indicated in the figure legends and the text. A significant difference was concluded at *P* < 0.05. Throughout all figures: \**P* < 0.05, \*\**P* < 0.01, \*\*\**P* < 0.001, and \*\*\*\**P* < 0.0001. Unless indicated otherwise all box plots show median, first and third quartiles, and whiskers represent minimum and maximum.

### Data availability

All data analyzed from published studies are referenced and publicly available under the following accession numbers: TCGA-SKCM dbGaP accession phs000178.v10.p8. Experimental data will be available from the corresponding author upon request.

## Acknowledgements

We thank members of our laboratory for comments on previous versions of the manuscript. We are grateful for assistance by Rockefeller University resource centers: Svetlana Mazel and staff of the flow cytometry resource center for assistance with flow cytometry, and Vaughn Francis and other veterinary staff of the Comparative Bioscience Center for animal husbandry and care. We thank the group of Marcus Bosenberg for generating the *Tyr::CreER;B-Raf^V600E/+^;Pten^lox/+^* mouse model and making it available through Jackson laboratories. We also express gratitude to Marcus Bosenberg for kindly providing the YUMM1.7 cell line. We thank the groups of Patrick M. Sullivan and Nobuyo N. Maeda for generating the human *APOE* targeted replacement mice and making them available through Taconic Biosciences. We are grateful to Elizabeth McMillan for help with whole exome sequencing analysis. We also thank Michael Szarek for statistical advice. This work was supported by a National Institutes of Health grant R01CA184804-01A2 (to S.F.T.). B.N.O. was supported by a Deutsche Forschungsgemeinschaft (DFG) postdoctoral fellowship (OS 498/1-1). K.T. and J.B. were supported by scholarships from the German National Academic Foundation. N.A. was supported by Medical Scientist Training Program grant T32GM007739 from the National Institutes of Health. B.T. was supported by the Lucy Lee Chiles Fellowship from the Hope Funds for Cancer Research. S.F.T. was supported by a Faculty Scholars grant from the Howard Hughes Medical Institute and the Reem-Kayden award.

## Author contributions

B.N.O. and S.F.T. conceived the study, supervised all research, and wrote the manuscript. B.N.O., N.A., K.T., J.B., and B.T. conducted experiments. B.N.O. analyzed whole exome sequencing and clinical data.

## Author information

S.F.T. is a co-founder, shareholder, and member of the scientific advisory board of Rgenix. S.F.T and B.N.O. are inventors on a US provisional patent application encompassing aspects of this work. Correspondence and requests for materials should be addressed to S.F.T.

## Extended Data

**Extended Data Fig. 1.**
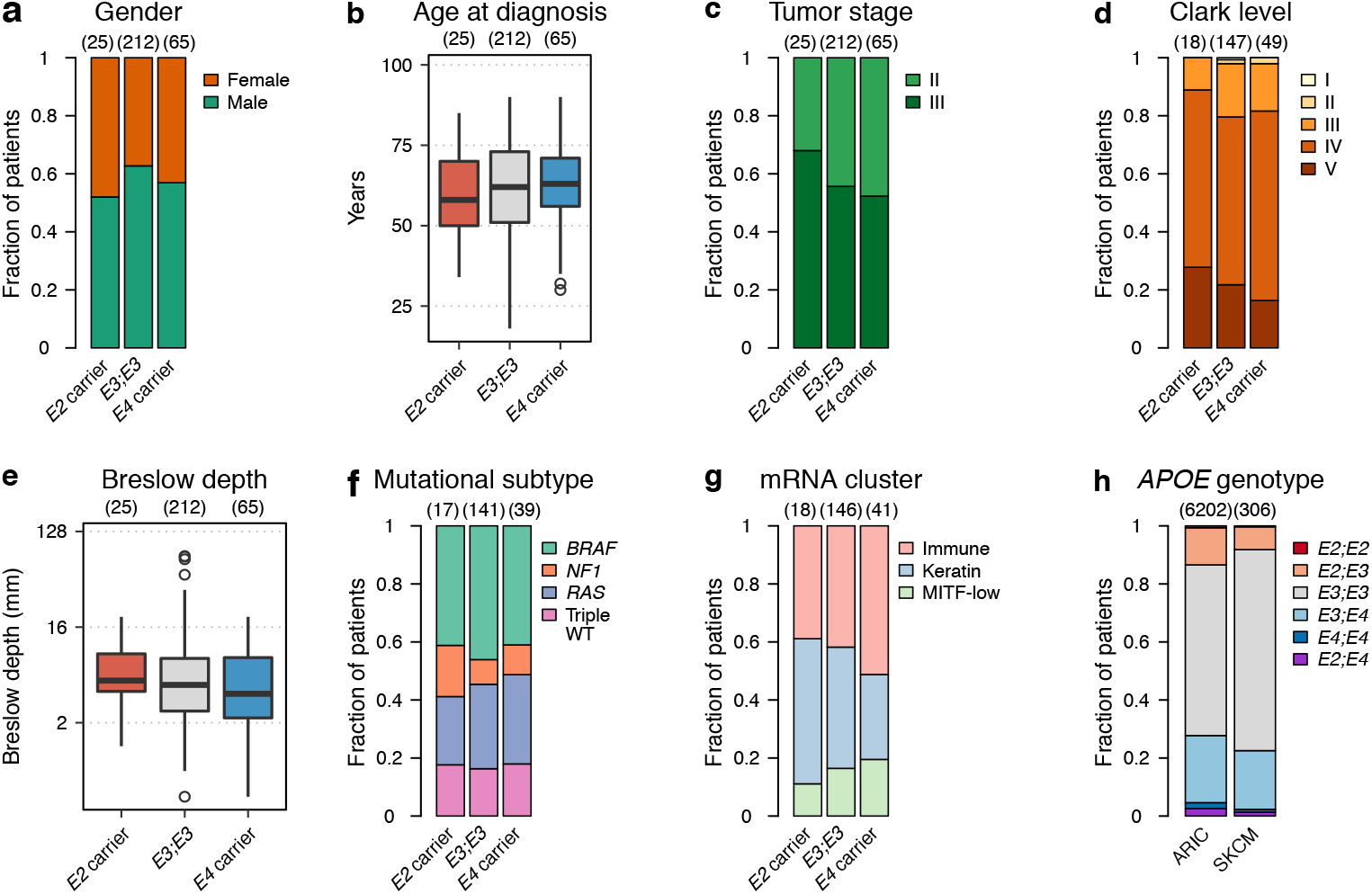
Clinical characteristics of stage II/III patients in the TCGA-SKCM cohort and bi-allelic *APOE* distribution. **a**, Gender proportions were not significantly different between *APOE* carrier groups (*P* = 0.46, χ^2^ test). **b**, Age at diagnosis was not significantly different between *APOE* carrier groups (*P* = 0.45, Kruskal-Wallis rank sum test). **c**, Tumor stage at diagnosis was not significantly different between *APOE* carrier groups (*P* = 0.4, χ^2^ test). **d**, Melanoma Clark level at diagnosis was not significantly different between *APOE* carrier groups (*P* = 0.95, χ^2^ test). **e**, Breslow depth was not significantly different between *APOE* carrier groups at diagnosis (*P* = 0.24, Kruskal-Wallis rank sum test). **f**, *APOE* carrier status was not significantly associated with tumor mutational subtype (*P* = 0.93, χ^2^ test). Proportion of *APOE* carrier status across molecular subtypes of the TCGA-SKCM cohort. **g**, *APOE* carrier status was not significantly associated with transcriptomic cluster (*P* = 0.55, χ^2^ test). Proportion of *APOE* carrier status across transcriptional clusters of the TCGA-SKCM cohort. **h**, *APOE* genotype distribution in the *Atherosclerosis Risk in Communities* (ARIC) cohort and stage II/III patients in the TCGA-SKCM study (*P* = 0.0066, χ^2^ test). Hinges of boxplots represent the first and third quartiles, whiskers extend to the smallest and largest value within 1.5 × interquartile ranges of the hinges and points represent outliers.

**Extended Data Fig. 2.**
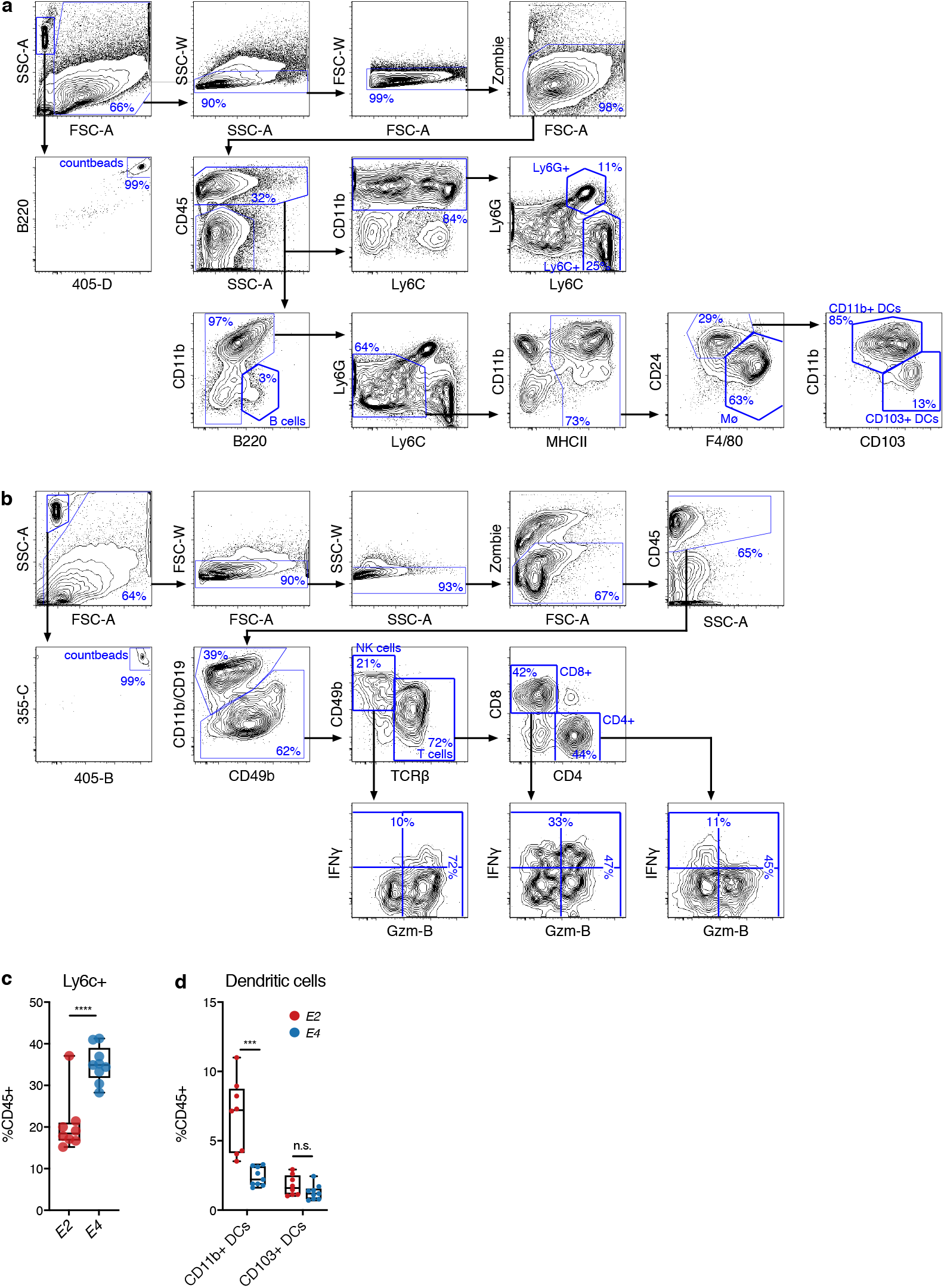
Immunoprofiling of the tumor microenvironment in *APOE2* versus *APOE4* mice. **a-b**, Representative flow cytometry plots demonstrating the gating strategy to identify major myeloid **(a)** and lymphoid **(b)** cell subsets in the tumor microenvironment. **c-d**, Proportion of Ly6C+ **(c)** and dendritic cell **(d)** subsets in the immune microenvironment of YUMM1.7 tumors in *APOE2* and *APOE4* mice (two-tailed t-test).

